# ORF Capture-Seq: a versatile method for targeted identification of full-length isoforms

**DOI:** 10.1101/604157

**Authors:** Gloria M. Sheynkman, Katharine S. Tuttle, Elizabeth Tseng, Jason G. Underwood, Liang Yu, Da Dong, Melissa L. Smith, Robert Sebra, Tong Hao, Michael A. Calderwood, David E. Hill, Marc Vidal

## Abstract

Most human protein-coding genes are expressed as multiple isoforms. This in turn greatly expands the functional repertoire of the encoded proteome. While at least one reliable open reading frame (ORF) model has been assigned for every gene, the majority of alternative isoforms remains uncharacterized experimentally. This is primarily due to: i) vast differences of overall levels between different isoforms expressed from common genes, and ii) the difficulty of obtaining contiguous full-length ORF sequences. Here, we present ORF Capture-Seq (OCS), a flexible and cost-effective method that addresses both challenges for targeted full-length isoform sequencing applications using collections of cloned ORFs as probes. As proof-of-concept, we show that an OCS pipeline focused on genes coding for transcription factors increases isoform detection by an order of magnitude, compared to unenriched sample. In short, OCS enables rapid discovery of isoforms from custom-selected genes and will allow mapping of the full set of human isoforms at reasonable cost.

## Introduction

Mechanisms that enable production of multiple isoforms from a single gene—alternative transcriptional start sites, splicing, and polyadenylation—contribute to expanding the functional capacity of the encoded proteome^1–3^. The full extent of this capacity is unknown, as we are currently unable to generate an accurate and comprehensive map of the human transcriptome^4^. Although advances in high-throughput sequencing have enabled mapping of local elements (e.g., individual splice junctions), how these elements combine to form full-length isoforms is largely unknown. Short read RNA-seq data from currently popular platforms (<250 bp, Illumina) fail to resolve such sequences^5, 6^. Consequently, a majority of annotated isoform models remain as predictions derived from partial transcripts, particularly for context-specific, disease-specific, or low abundance isoforms^4, 7^.

Long-read sequencing platforms—PacBio^8^, Oxford Nanopore^9^, and those based on adaptations to next generation sequencing that produce synthetic long reads such as 10X and Moleculo^10^ sequencing—can return unambiguous, full-length isoform sequences that fully resolve transcriptome complexity. However, in comparison to short-read sequencing platforms, these methods suffer from lower sampling sensitivity and can miss low (<10 copies/cell) to moderate (10-50 copies/cell) abundance transcripts. Transcripts at these levels include many disease-associated or important regulatory proteins (e.g., transcription factors and kinases). The sensitivity problem is exacerbated by the wide dynamic range of the human transcriptome across at least six orders of magnitude^11^, causing an inordinate amount of sequencing effort to be used for detecting the most abundant isoforms. Therefore, most transcripts of lower abundance have not been sequenced satisfactorily due to this biased sampling.

An established solution to increase detectability of isoforms is targeted sequencing, which involves enriching for desired transcripts from a population of RNA or cDNA molecules. For this purpose, DNA or RNA hybridization-based enrichment followed by high-throughput sequencing is particularly efficient, robust, and cost-effective^12^. These approaches were initially developed for targeted sequencing of protein-coding regions of genomic DNA (whole exome sequencing)^13^ and RNA fragments from short-read RNA-seq experiments (e.g., CaptureSeq)^14–18^. Such approaches have been adapted for targeted sequencing of long genomic fragments (>2kb) or full-length cDNA molecules^19–24^. In two notable studies described recently, complex pools of biotinylated oligos were used to enrich for and sequence thousands of protein-coding and lncRNA targets, leading to considerable gains in isoform detection and new insights about the nature of transcriptomic complexity^25, 26^.

The success of these targeted full-length sequencing methods hints at the potential of using this approach in a more general framework. These methods employed a single probe panel with many targets. This model has worked well for exome sequencing, in which a single probe set designed against all protein-coding exons yields high coverage of all DNA targets, each of which are present at identical concentrations (2 copies/cell). However, such complete coverage is challenging to attain from targeted transcriptome sequencing. Transcriptomes are heterogeneous, with a wide dynamic range and wide variation in composition (set of genes expressed), depending on cellular or disease context. Therefore, a single probe set will enable increased sequencing of isoforms from genes of interest, but the expression patterns within the genes of interest will still be skewed, reducing coverage. Therefore, what is needed is a flexible strategy to generate with ease and low-cost multiple, distinct probe sets that match the particular transcriptome context, specifically, to enable facile detection of isoforms from any set of genes from any set of samples.

We developed “ORF Capture-Seq” (OCS), a pipeline that provides a generalizable method for direct synthesis of biotinylated capture oligos from existing or newly designed ORF clones followed by targeted enrichment and sequencing of full-length cDNAs to discover isoforms in any tissue of interest. We emphasize that the unique combination of low cost, time, ease, and versatility (any pool of ORFs/clones, up to thousands at-a-time) of the method offers the experimental flexibility needed to rapidly characterize any desired subset of the transcriptome. Using reagents and instruments available in most molecular biology labs, a user can synthesize probes from one or a set of amplicons or clones in less than 24 hours. We envision this method will be of broad utility in many applications, from single-gene studies to system-scale applications seeking to characterize whole transcriptomes. Here, we compare OCS probes against a commercial standard, benchmark the method using spike-in standards, and apply it towards characterization of novel isoforms of ∼800 human transcription factors (TFs).

## Results

### OCS method for flexible targeted sequencing

The OCS pipeline begins with flexible and straightforward synthesis of biotinylated capture probes (Fig. 1a). PCR is performed on any number of pooled templates (e.g., plasmids, amplicons) using universal primers in the presence of biotin-dUTP. The resulting pool of biotinylated PCR products, with biotin-dUMP incorporated throughout, are randomly fragmented to an average size of ∼150 base pairs to generate a set of overlapping fragments from each PCR amplicon. After purification and removal of PCR primers and unincorporated nucleotides, the resulting OCS probe set is used for hybridization-based capture of target nucleotide sequences.

**Fig. 1.**
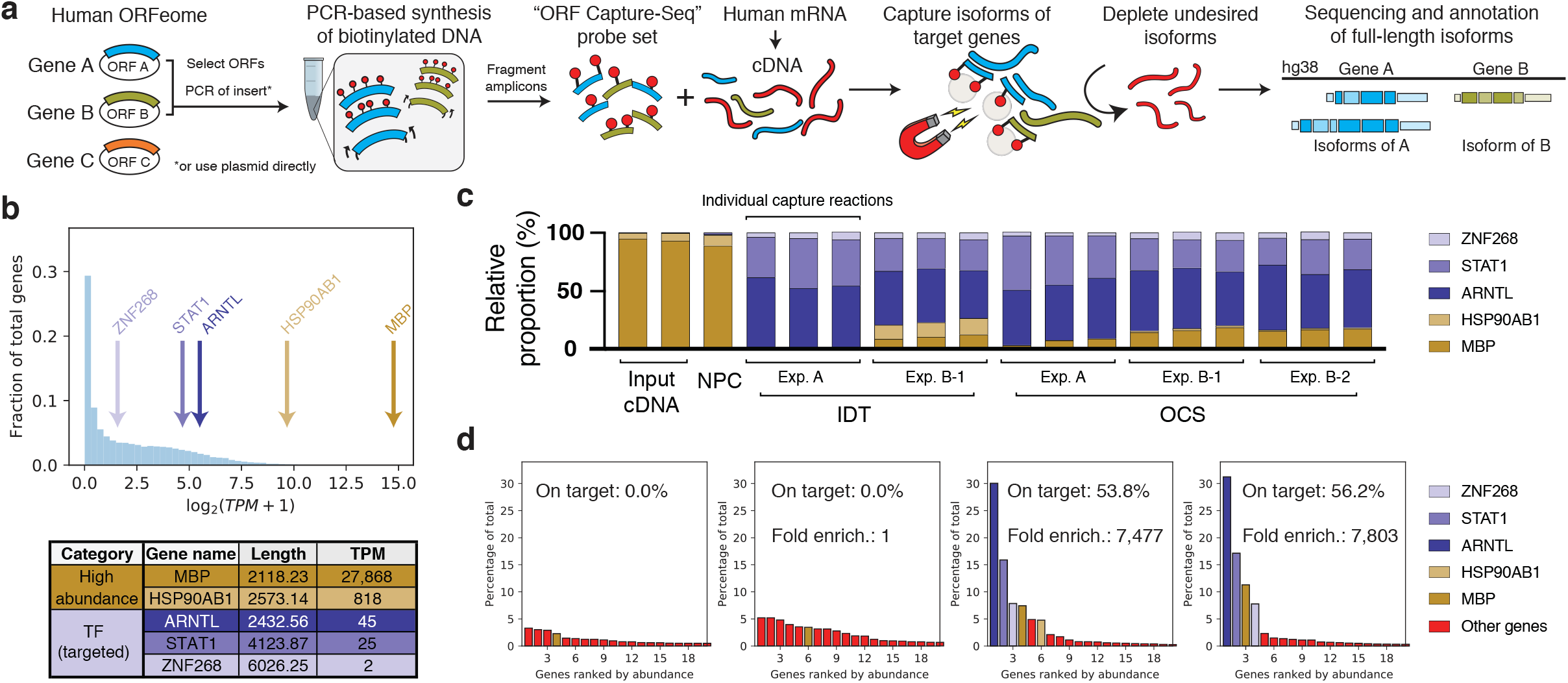
ORF Capture-Seq (OCS) method for accelerated discovery of full-length isoforms. **a** Schematic of the OCS method. ORF clones of target genes are pooled and used as templates for a Biotin-dUTP-labeling PCR reaction, creating randomly biotinylated amplicons which are fragmented to generate a probe set. In this study, PCR-based amplicons derived from the clones were used as template. These OCS probes can be used in targeted sequencing applications, such as enrichment of full-length cDNA for sequencing on the PacBio platform. **b** Transcriptional abundances in human brain cDNA. These values were used as the basis for selecting three low to moderate abundance transcription factors (TFs) as target genes (purple labels) and two high abundance genes (yellow labels) as background controls. Length is an average of all transcripts annotated for each gene (GENCODE v22). TPM values were obtained from processing Illumina sequencing data (Methods). TPM, transcripts per million. **c** Comparison of IDT vs OCS-based target enrichment. Each bar shows the relative proportion of cDNA from target (purple) versus background (yellow) genes as quantified by qPCR (average of two technical replicates). NPC, no probe control. **d** Plots of individual gene expression, ranked in descending abundance, as quantified by Illumina sequencing and Kallisto (Methods). Each bar represents one gene and the 20 most abundant genes are shown. Bars are color coded as background controls (yellow), target genes (purple), and all remaining genes that were not targeted (red). On-target percentages are the fraction of transcriptional abundance corresponding to the three targeted TFs (ARNTL, STAT1, ZNF268), in each capturant. Fold enrichment is computed by dividing percentage of targets in capturant by the percentage in the input.

We demonstrate the application of OCS for enrichment and sequencing of full-length isoforms from protein-coding genes (Fig. 1a). Probes are derived from one or more ORFs per gene. Though each ORF represents just one isoform of a gene, the corresponding probes are expected to capture the family of isoforms from that gene, due to the high sequence overlap between isoforms of the same gene; probes need only target a portion of a full-length cDNA for enrichment^16^. Thus, we capitalized on the availability of the human “ORFeome” collection at the Center for Cancer Systems Biology (CCSB) at the Dana-Farber Cancer Institute, a resource of “ready-to-express” and freely available ORF clones for ∼17.5K of the ∼20K protein-coding genes in human^27^. From this resource, we created customized pools of ORF clones to use as templates (corresponding to the genes of interest to target for enrichment) for biotinylated PCR. All clones share a common vector backbone, enabling production of any amplicon from universal primers. Hundreds to thousands of ORF clones may be pooled and processed in this manner so that complex and customized probe sets can be generated with relative ease.

### OCS probes perform comparably to a commercial standard

We first established that OCS probes are comparable to commercially synthesized biotinylated probes, in terms of enrichment efficiency.

For benchmarking, we selected three low abundance, brain-expressed human TF genes (*ARNTL*, *STAT1*, and *ZNF268*). The enrichment of these TFs were compared against two high abundance housekeeping genes (*MBP* and *HSP90A1)* to serve as a measure of off-target binding (Fig. 1b). For each of the three TFs, OCS and commercial probes were synthesized (Supplemental Fig. 1a). OCS probes were sequenced on an Illumina MiSeq, confirming an even distribution of probe coverage (estimated ∼150X tiling density) and high purity (Supplementary Fig. 1b-d, Methods). Commercially available probes were synthesized as 5’ biotinylated 120-mers with a ∼1X tiling density against both the forward and reverse strands (Supplementary Fig. 1a, Supplementary Table 1, Methods). Probes were synthesized by Integrated DNA Technologies (IDT) and hereafter are referred to as “IDT probes”.

We compared the ability of OCS and IDT probes to enrich for transcripts corresponding to the three TF genes from human brain cDNA—in technical triplicate and with the entire experiment repeated on a different day (experiment A-1 and B-1 in Fig. 1c, Methods). A capture reaction employing OCS probes from a second independent synthesis was also performed (experiment B-2 in Fig. 1c). In each capturant (i.e., post-capture cDNA), concentrations of transcripts corresponding to the three TF target genes and the two non-target background genes were measured using qPCR (Fig. 1c). We found that the major cDNA population corresponded to the three target TFs (∼80% on-target). Importantly, the on-target enrichment rates, defined as the fold increase in relative abundances of the TFs, were statistically indistinguishable between OCS and IDT probes (Supplementary Fig. 1e), given the intra-assay (CV=2.2%) and inter-assay (CV=10.3%) variability. Finally, a subset of the capturants were subjected to sequencing on an Illumina MiSeq (∼50K 150 paired end reads per sample, Methods), providing estimates of on-target rates of 54% for OCS and 56% for IDT (Fig. 1d). Thus, OCS probes perform comparably to a commercial standard.

The OCS probes were derived from PCR inserts from Gateway clones, in which each ORF is flanked by attB sites and ∼100 bp of vector backbone^27^. A possible concern is that probe sequences arising from the vector backbone can cause non-specific binding. To investigate this, we compared background binding profiles derived from OCS versus IDT capture experiments. The profiles are displayed as enrichment of each transcript as a function of initial abundance, because higher abundance transcripts have been observed to non-specifically bind to the beads to a greater extent than low abundance transcripts (Supplemental Fig. 1f, see “No Probe Control”). We found no systematic biases using either probe type.

### Analytical benchmarking of OCS using spike-in standards

Next, we benchmarked the analytical performance of OCS by employing External RNA Controls Consortium (ERCC) standards, which are 92 synthetic ORFs of concentrations spanning 10-orders of magnitude^28^.

To assess specificity and reproducibility, we measured enrichment of a subset of ERCC ORFs in human reference RNA (Fig. 2a). OCS probes were synthesized for the 64 ERCC ORFs in the lowest concentration bracket (“ERCC64”, Supplementary Fig. 2a, Supplementary Table 2, Methods). The remaining 28 ORFs of highest concentration were not targeted and served as background controls. The method of probe synthesis used was analogous to the human ORFeome-based synthesis described above, except that plasmids containing ERCC ORFs served as templates (Methods). ERCC RNA standards were spiked into 1 µg Universal Human Reference RNA (UHRR) at dilution levels of 1:10 (i.e., 10X dilution), 1:80, and 1:5120, in technical duplicate (Supplementary Fig. 2b, Methods). ERCC64 was used to enrich for target transcripts. Input and capturant cDNA were sequenced on an Illumina MiSeq and abundance values estimated using Kallisto^29^, in which a subsampling of 100K reads were subjected to analysis to allow for comparison of ORF detection at comparable sequencing depths.

**Fig. 2.**
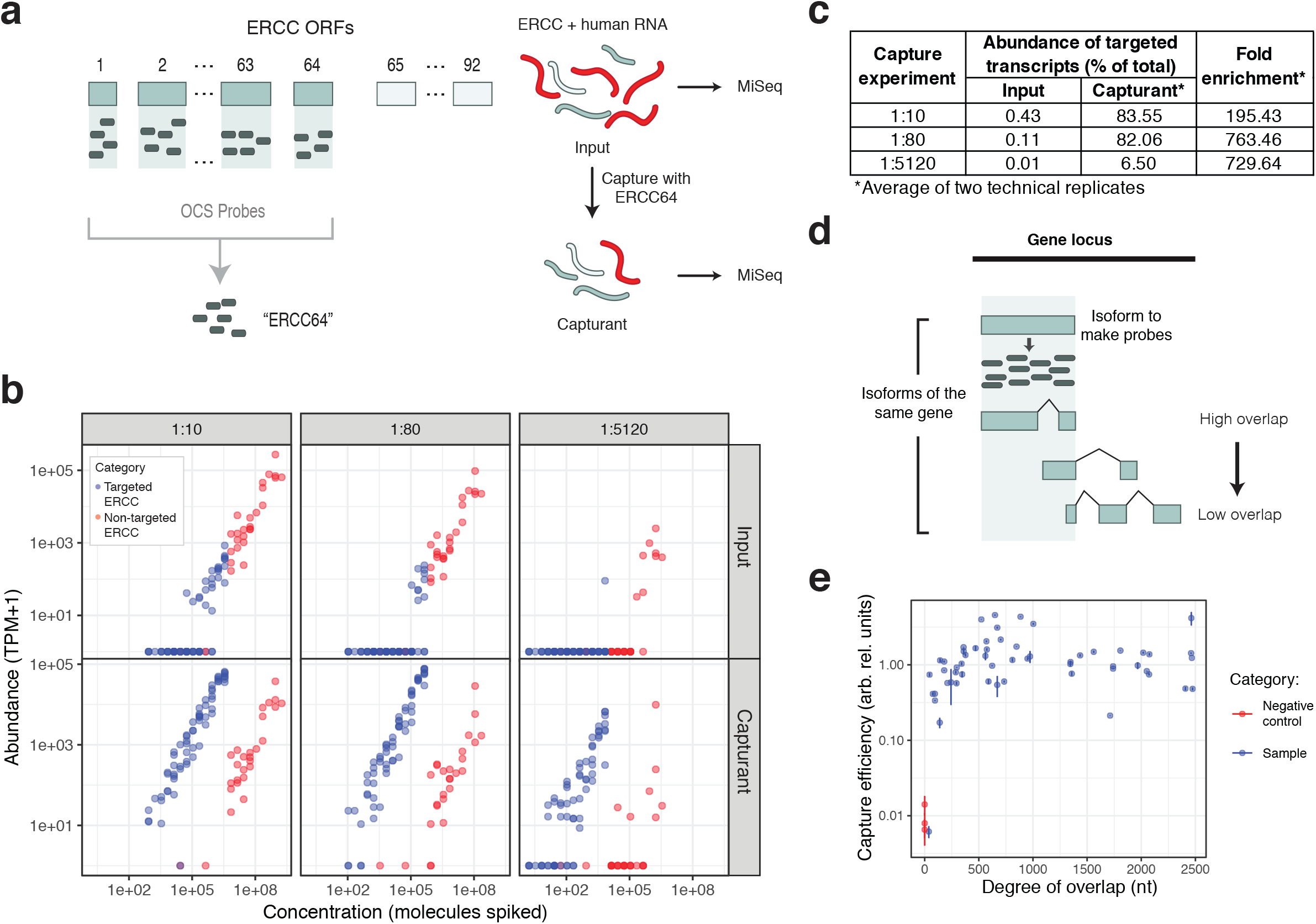
Benchmarking OCS analytical performance. **a** Schematic of benchmarking experiment using ERCC standards. **b** Plots showing enrichment of ERCC targets. The x-axis represents the nominal concentration of ERCC RNAs spiked into the starting pool of RNA (input) and the y-axis represents estimated abundance of each ORF in the input cDNA (top row) or capturant (bottom row). Each point represents a distinct ERCC standard (92 total) which was targeted (blue) or not targeted (red). **c** Table of summary statistics related to capture efficiencies for ERCC capture reactions. **d** Schematic of probe synthesis using the SIRV system. **e** Plot showing the relationship between enrichment efficiencies at different isoform overlaps. The isoform overlap represents the absolute number of nucleotides overlapping between the 1) template isoform used to generate probes, and 2) target isoform present in the sample. Negative controls are non-overlapping isoforms. Capture efficiencies were computed by dividing read depth of each SIRV (isoform) in capturant by the read depth in input cDNA.

The ERCC capture experiments demonstrated excellent specificity with uniformly elevated abundances across all 64 targeted ORFs (Fig. 2b). We computed overall on-target rates by computing the total abundance, in transcripts per million (TPM), arising from ORFs targeted for enrichment. Fold enrichment was calculated by dividing on-target percentages found in the capturant by those found in the input. For the 1:10 and 1:80 capture, on-target rates were above 80% with fold enrichments of 195- and 763-fold (Fig. 2c, Supplementary Fig. 2c). For the 1:5120 capture, on-target rates were lower (∼6%) but enrichment was high (730-fold). The relative abundances of targeted ERCCs remained linear post-capture indicating potential for deriving quantitative information from captured material, given that probes are in excess concentrations compared to the target cDNA (as previously observed^30^) (Supplementary Fig. 2d). Furthermore, the extent of enrichment was not significantly biased by properties such as starting concentration, GC content, ORF length, and probe representation (Supplementary Fig. 2e). Technical reproducibility was also high, with Pearson’s r-squared above 97% for all technical replicates (Supplementary Fig. 2f).

### OCS can enrich for the family of isoforms using one representative isoform

The use of OCS for isoform discovery relies on the assumption that probes derived from a single isoform are sufficient to enrich for the family of isoforms of the same gene. Indeed, there is typically sufficient overlap between any given isoform sequence and all other isoforms of that gene. However, to address the concern that there could be lower capture efficiency due to low sequence overlap, we measured the relationship between overlap and capture efficiency.

For this purpose, we used Spike-in RNA Variant (SIRV) standards (Lexogen), 69 synthetic isoforms with highly complex splicing patterns from seven genes (Fig. 2d)^31^. We synthesized OCS probes from one representative SIRV isoform per gene (“SIRV7”, Supplementary Fig. 2g-h, Supplementary Table 3, Methods) and used this probe set to attempt enrichment for all SIRV isoforms that were spiked into UHRR (Supplementary Fig. 2i). We found no appreciable difference in capture efficiency when sequence overlap ranged between 50 and 2500 nt (Fig. 2e). In extreme cases in which overlap was lower than 50 nt, capture efficiency sharply declined to the level of negative controls.

To estimate the isoform space covered in the application of this method for protein-coding human genes, we calculated the degree of overlap between the principal isoform, as defined in GENCODE^32^ (version 29), and all other annotated isoforms of the same gene (Supplementary Fig. 2j, Methods). The overlap was calculated by taking the intersection of genomic ranges. We found that 99.7% of all isoforms are potentially captured by OCS probes designed against the principal isoform.

### Applying OCS to characterize human TF isoforms

Alternative transcriptional start sites, splicing, and/or polyadenylation can modulate the activity of TFs by altering sequences corresponding to DNA binding, co-factor binding, and other properties such as availability of phosphorylation sites^33, 34^. Despite being heavily studied, many TF isoforms remain uncharacterized due to low abundance (<10 copies per cell), complex splicing patterns, or expression in cell-, tissue-, or disease-specific contexts. We applied OCS to characterize alternative isoforms of human TFs.

We tested the extent to which the number of targeted TF genes could be multiplexed in OCS experiments, beyond the numbers of genes we tested heretofore (∼100 genes). We sought to explore the relationship between the number of genes and sensitivity. First, we synthesized OCS probe sets corresponding to 2, 12, 88, and 682 TF genes and used each set to enrich for TF isoforms from human cDNA derived from cerebral cortex (Supplementary Table 4, Methods). The probe sets were subjected to sequencing and the relative representation of each TF was inferred from the relative abundances of the corresponding probes, confirming high purity and coverage (Supplementary Fig. 4a, Supplementary Table 5-6, Methods). Capturants from the experiments were sequenced on an Illumina Next-Seq and on a PacBio RSII (Methods). With increasing OCS probe set complexity (i.e., number of genes represented), we found that overall capture efficiency was high (Fig. 3a) and that a higher absolute number of genes and isoforms were identified overall (Fig. 3b). However, increased breadth came at a cost of decreased depth—a lower fraction of genes was detected and genes in larger probe sets returned fewer isoforms per gene on average (Fig. 3c-d). This observed trend was confirmed by saturation-discovery curve analysis (Supplementary Fig. 3a).

**Fig. 3.**
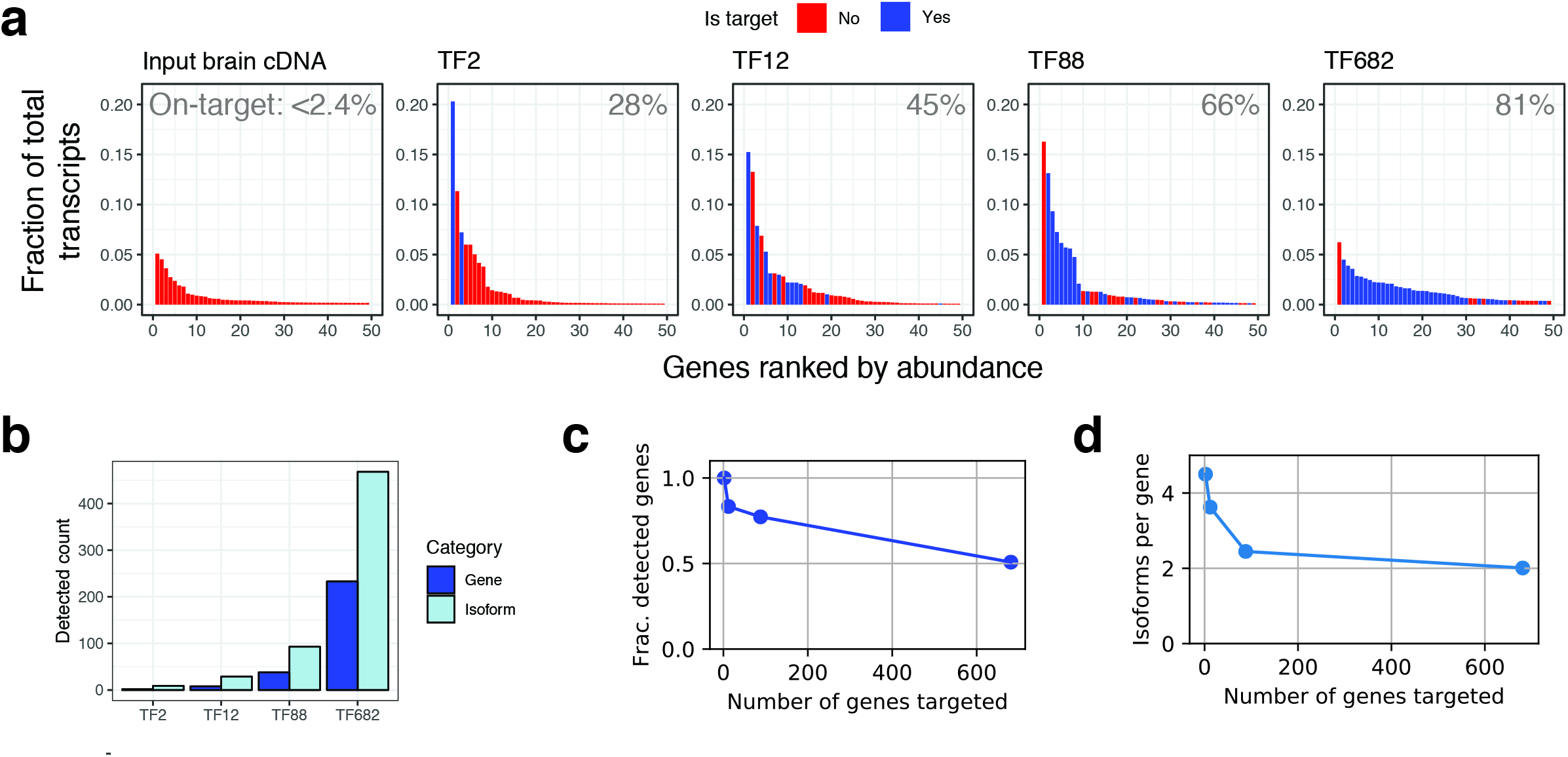
Multiplexing parameters for enrichment of human transcription factors. **a** Rank abundance bar plots for unenriched (input) and enriched cDNA. Data is shown for 1) input brain cDNA, and 2) the series of enriched cDNAs from performing capture reactions using probe sets with increasing number of TF genes. Only the top 50 most abundantly expressed genes, calculated per sample, are shown. Each bar corresponds to a single gene, colored by whether that gene is targeted (blue) or not targeted (red). Fraction of total transcripts was calculated by dividing the transcript abundance (TPM) of all transcripts from a gene by the total transcript abundances for the sample (Methods). On-target rates, as calculated for the entire sample, are displayed on the upper right-hand position of the plots. **b** Bar plot showing the absolute number of targeted genes (dark blue) and isoforms (light blue) detected from each capture reaction. **c** Plot showing the relationship between number of genes multiplexed and the fraction of genes for which there was a detected full-length read. Frac., fraction. **d** As in **c** except shows the decrease in isoforms per targeted gene, on average, for each experiment.

We applied OCS towards the discovery of TF isoforms from a diverse set of human tissues. Based on our goal of increased breadth, we decided to employ a large probe set comprising 763 TFs (“TF763”) to enrich from a pool of 7 barcoded human tissue cDNA libraries (Supplementary Fig. 4a, Supplementary Table 7-8, Methods). Both the unenriched input and enriched capturant cDNA was subjected to PacBio sequencing on 1M SMRTcells on the PacBio Sequel system. Raw data was processed using the Iso-Seq v3 bioinformatic pipeline^35^ followed by SQANTI for isoform annotations^36^. Post-processing resulted in a total of 118,872 and 476,589 full-length reads (circular census reads, or CCS), originating from the input and capturant, respectively. In the capturant, the on-target rate was ∼60%, with a majority of the top ranked genes arising from target TFs (Fig. 4a). To compare efficiency of enrichment, ∼100K full-length reads were each sampled from the input and capturant datasets and sequencing statistics were calculated. We found that the number of genes, isoforms, and full-length reads increased 2-, 7-, and 43-fold after enrichment respectively, emphasizing the need for enrichment of some sort to progress towards fully sampling the isoform space (Fig. 4b).

**Fig. 4.**
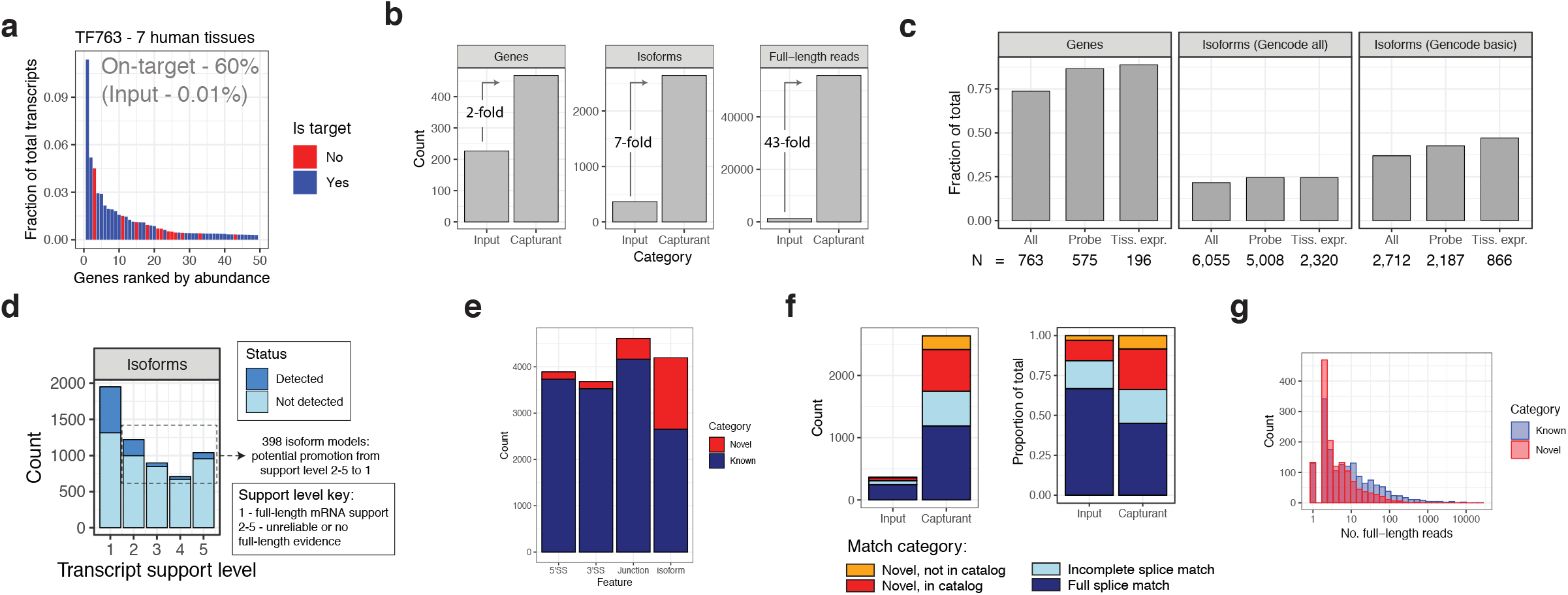
Full-length transcription factor isoforms across diverse human tissues. **a** Rank abundance bar plot for cDNA enriched for human TFs. Data for the top 50 most abundantly expressed genes, are shown. Each bar corresponds to a single gene, colored by whether that gene is targeted (blue) and not targeted (red). Abundances as shown on the y-axis are computed by dividing the number of full-length reads mapped to the gene by the total number of full-length reads. On-target rates for input (rank plot not shown) and capturant (associated with this rank plot) are shown. **b** Bar plot showing gains in coverage of target genes upon enrichment. Increases in number of genes, isoforms, and full-length reads are shown. Data used to generate these numbers used an equal number of full-length raw reads that were subsampled from the un-enriched (input) or enriched (capturant) cDNA. **c** Plot of the fraction of all GENCODE genes and transcripts detected in the capturant. A gene is considered detected if at least one full-length read is detected for that locus. Isoforms are considered detected if the full set of junctions are identical between the GENCODE-annotated and sequenced transcript. The fraction detected was also computed for sets of genes for which there was higher probe set representation of the gene (1 TPM or higher, label: Probe) and genes for which there was evidence of expression in the tissues interrogated (10 TPM or higher, label: Tiss. Expr.). The number of genes or isoforms involving the original or subset of genes are shown at the base of each bar. Tiss. Expr., Tissue expression. **d** Full-length sequence data applied towards validation of existing gene models. Plot shows for each transcript support level the number of isoforms annotated in GENCODE, for the genes targeted in the TF enrichment experiment (TF763). Fraction of all isoforms that exactly match, and thus is validated by, a full-length sequenced transcript, is shown in dark blue. **e** Fraction of novel splice sites, junctions, and full-length isoforms in the TF enrichment experiment. Unique splice sites and junctions are only counted once. The 5’ splice site corresponds to the splice donor and the 3’ splice site corresponds to the splice acceptor. SS, splice site. **f** Stacked bar plots of the proportions of known and novel isoforms. Both absolute (left plot) and normalized (right plot) versions are shown, for comparison. Known isoforms are further divided by completeness. Novel isoforms are further divided by whether all splice sites are found in GENCODE (novel in catalog) or if the isoform contains a novel splice site (novel not in catalog). Match categories are based on the isoform annotation tool SQANTI. **g** Distribution of full-length read depth for known and novel isoforms.

We analyzed the extent to which we recovered full-length isoforms that are annotated in GENCODE (Fig. 4c, Supplementary Table 9). Overall, the recovery of GENCODE genes was 74%, increasing to 86% or higher when considering only genes which were well-represented in the probe set (1 TPM or higher in the set of biotinylated probes, TF763), or genes well expressed in the 7 tissues we subjected to analysis (10 TPM or higher). The recovery of GENCODE isoforms was 22%, also increasing when accounting for probe representation or tissue expression. Isoform recovery was yet higher (37%) for GENCODE high-confidence isoforms. As expected, isoforms detected in the few samples we interrogated represent a fraction of all possible isoforms expressed in human samples. Indeed, isoform recovery will not be 100%, as GENCODE models are derived from sequencing data collected from hundreds of diverse tissue and cell types, whereas this study was restricted to seven normal human tissues.

In GENCODE annotations, every isoform is tagged with a transcript support level (TSL) 1-5, denoting the extent of experimental information underlying each isoform model. Isoforms with a TSL of 1 contain high-quality, full-length mRNA sequence support. Isoforms with a TSL of 5 are computational predictions with no experimental support. The full-length sequencing data provided confirmation of 398 GENCODE isoforms of TSL 2-5, which includes 85 computationally predicted isoforms (TSL 5) (Fig. 4d, Supplementary Table 10). Thus, OCS is an invaluable tool for confirming isoform models in gene annotations.

OCS-based enrichment enables significant increases in detection of annotated isoforms as well as discovery of novel isoforms. Given the high number of novel isoforms routinely detected in long read sequencing studies, an open question in the field is how to best assess the quality of the isoforms. Few studies have systematically investigated the sources and frequency of artifacts, but one of the most comprehensive assessments to date is work from Tardaguila and colleagues^36^, in which they evaluated intrinsic and sequence-related properties that contribute to isoform artifacts. They found that non-canonical or RT-template switching junctions underlie poor quality novel isoforms, but that experimental validation using an orthogonal approach effectively detected and removed these events.

Because of the demonstrated value of orthogonal sequencing data to validate novel isoforms, we subjected the original RNA from the samples analyzed in this study to dication-catalyzed RNA fragmentation and random hexamer priming to generate cDNA fragments which were subjected to Illumina sequencing (Methods). The Illumina reads were used to validate isoforms detected using the OCS approach; we required each junction in a novel isoform to be supported by a minimum of three Illumina reads.

This filtering approach resulted in a population of novel isoforms with quality features (e.g., non-canonical junction rate), intrinsic sequencing properties (e.g., number of predicted RT artifacts), and functional genomics evidence (e.g., overlap of 5’ end with CAGE peaks) that are indistinguishable from the high-quality known isoforms that match GENCODE annotations (Supplemental Fig. 4b-j). All subsequent analyses involved the orthogonally-validated novel isoforms (Supplementary Table 11).

Using the high-quality set, isoforms and their specific junctions and splice sites were evaluated for their novelty, defined here as not being in GENCODE v29 (Fig. 4e). Approximately 4% and 10% of all distinct splice sites and junctions were novel, respectively, but a much higher fraction of isoforms, 37%, were novel. This can be explained by the fact that a single local event (e.g., one novel splice site) leads to an entirely distinct transcript, in terms of the full-length sequence. Overall, the total number of isoforms detected dramatically increased upon enrichment and at the same time the relative fraction of novel isoforms increased in proportion (Fig. 4f). A substantial proportion of the reads arose from novel isoforms, though the number of full-length read counts were slightly lower for novel isoforms as compared to known isoforms (Fig. 4g). Overall, 1,528 distinct novel isoform sequences were detected.

## Discussion

Eukaryotic transcriptomes remain unresolved at full-length resolution, and the full extent of transcript diversity is unknown. Recently, targeted full-length sequencing methods have characterized focused subsets of the transcriptome with great depth and accuracy. Here, we establish that ORF Capture-Seq is a versatile method to synthesize biotinylated probes that can be used in targeted full-length sequencing studies. Applying OCS to detect TF isoforms in human tissues, we found a preponderance of novel isoforms, detecting over a thousand novel TF isoforms. OCS can be applied for isoform discovery for other classes of proteins, such as kinases or G protein-coupled receptors, or proteins involved in a biological pathway of interest or implicated in genomic studies of a human disease.

OCS enables direct characterization of isoform sequences without prior information about transcript boundaries or exons, unlike with RACE or PCR-based sequencing. This facilitates discoveries of the full array of transcripts in specific conditions, diseases, cell-types, and single cells. Such information could guide high-throughput cloning efforts^37^. OCS can help define full-length isoform sequences from genes exhibiting differential splicing at the local level, using programs like Leafcutter^38^. It also provides opportunities for increased accuracy in isoform quantification workflows. Knowledge of isoforms expressed in a sample, as informed by long-read data, can serve as the “scaffold” (i.e., gene models) upon which short reads rely to estimate isoform abundances^36, 39^.

The ease of making probes using the OCS method opens doors for novel strategies in capture experiments. For example, a series of captures can be designed in an iterative manner, in which the initial capture returns the first “batch” of detected genes, and subsequent captures use probe sets that include only genes that failed to return isoforms in the first round. Alternatively, multiple gene panels may be created, stratified by endogenous abundance of genes (e.g., separate probe sets for low and high abundance transcripts) or priority of disease genes (e.g., low/high confidence of association). Furthermore, the representation of genes within a probe set is customizable to the greatest extent. For example, the concentration of individual ORFs may be titrated based on a number of factors, such as priority or endogenous abundance in the sample, so as to normalize the proportions of genes in the capturant, allowing for greater sequencing coverage.

Some limitations of OCS remain. For example, since the method relies on PCR to synthesize probes, one limitation is the lengths of ORF clones that can be used as template; probe synthesis and full-length sequencing for transcripts above 5-6 kb was challenging. For longer genes, other mechanisms to generate biotinylated probes may be required (e.g., 5’ biotinylated directed^40^ or random primers, ligation^41, 42^, or nick translation^43–45^). Alternatively, we generated probes from individual segments of an ultra-long ORF clone (unpublished). Another limitation is related to the cDNA library itself. The use of whole-cDNA library PCR may also generate artifacts during the processing steps, so each isoform required orthogonal validation from Illumina sequencing data.

In conclusion, OCS is a highly generalizable strategy to synthesize probes for use in full-length capture experiments. Though we demonstrated OCS as applied towards characterization of isoforms from protein-coding genes, it can be adapted for use in capture and characterization of different types of genetic and post-transcriptional variants, such as genetic variations, segmental duplications, or lncRNAs. It is also possible that ORFs from one species could be used to enrich for isoforms from another species, given high sequence conservation of protein-coding regions (e.g., human ORFs to enrich for mouse isoforms). We envision this approach will be of broad utility for application within both basic research and the clinical and diagnostic fields^46^.

## Online methods

### ORF Capture-Seq probe synthesis

*ARNTL, STAT1, ZNF268 (three TFs)*

#### Generating ORF amplicons

The hORFeome contains one representative ORF, in the form of a Gateway clone, for ∼17,500 of the ∼20,000 human protein-coding genes^27^. These ORF clones are available as bacterial (DH5alpha) culture glycerol stocks. Bacterial stocks corresponding to the three TFs were cherry-picked from the hORFeome. Using ∼1µl of culture as template, the ORF inserts were PCR amplified with Platinum Taq polymerase (Invitrogen) using M13 primers with the following sequences:

> M13_FOR: CCCAGTCACGACGTTGTAAAACG
>
> M13_REV: GTAACATCAGAGATTTTGAGACAC

PCR was performed for 35 cycles, with each cycle consisting of denaturation at 94°C for 30 seconds, annealing at 57°C for 30 seconds, and extension at 72°C for 5 minutes. The final extension was for 15 minutes.

PCR products were analyzed via agarose gel electrophoresis to confirm that amplicons were of the expected size. Products were purified using Agencourt AMPure XP beads (Beckman Coulter) and quantified using the Qubit dsDNA HS assay (Thermo Fisher Scientific).

#### Biotin-labeling PCR

Both control and Biotin-dUTP PCRs were performed for each TF. Using as template 1 ng of the ORF amplicon, biotin-spiked PCR was done using Taq polymerase (NEB). The dNTP mixture was modified so that a third of the dTTPs were substituted with Biotin-16-Aminoallyl-2’-dUTP (Trilink), referred to heretofore as Biotin-dUTP. The program was run for 30 cycles, with each cycle consisting of denaturation at 95°C for 15 seconds, annealing at 57°C for 30 seconds, and extension at 68°C for 5 minutes. The final extension was for 10 minutes.

#### Fragmentation of amplicons

Control and biotinylated amplicons were run on an agarose gel to confirm successful PCR. Products were transferred to Covaris AFA FiberCrimp Cap microTUBEs and fragmented on a Covaris E220 sonicator to size distribution of ∼150 bp. The sonication method parameters are as follows: peak power of 175W, duty cycle of 10%, 200 cycles per burst, and duration of 480 seconds. Fragmented DNA was purified using SPRISelect beads (Beckman Coulter) using a 1:0.6 ratio of sample to beads to remove high mass fragments above ∼300bp. Concentration of fragments were measured with the Qubit dsDNA HS assay (Thermo Fisher Scientific).

#### Mixing of probe sets

An equi-mass mixture of the three TFs probes was prepared. The final concentration of the probe set was adjusted to 0.5 ng/µl.

For the following gene sets, OCS probes were synthesized using the same protocol as for the three TFs, with exceptions described in each section.

##### ERCC64

The External RNA Controls Consortium, or ERCC, has created a collection of 92 synthetic ORF sequences, in the form of plasmids, from which RNA standards have been prepared by various vendors. We obtained the ERCC DNA Sequence Library for External RNA Controls (SRM 2374, NIST), a collection of all ERCC ORFs in the pT7T318 vector.

ERCC ORF inserts were PCR amplified with hot-start KOD polymerase (Invitrogen) using M13 primers, sequences below:

> M13_canon_FOR: GTAAAACGACGGCCAGT
>
> M13_canon_REV: CAGGAAACAGCTATGAC

PCR was performed for 18 cycles, with each cycle consisting of denaturation at 94°C for 30 seconds, annealing at 55°C for 30 seconds, and extension at 72°C for 5 minutes. The final extension was for 15 minutes.

An amplicon pool was prepared for 64 ERCC ORFs (Supplementary Table 2). To make the ERCC64 amplicon pool, PCR reactions were pooled with adjustment based on length of the ORF, where higher volumes were used for longer ORFs. Using the pool of amplicons as template, a single Biotin-labeling PCR was done.

##### SIRV7

The Spike-in RNA Variant Control Mixes, or SIRV Mixes (Lexogen), are 69 synthetic transcripts from seven genes which mimic the highly complex splicing patterns found in the human transcriptome^31^. We obtained PCR products of SIRV constructs corresponding to SIRV101, SIRV201, SIRV301, SIRV403, SIRV510, SIRV601, and SIRV701. ORF-specific primers were used for the Biotin-labeling PCR and were designed to anchor the ATG-start and just upstream of the stop codon. All SIRV primer sequences used may be found in Supplementary Table 3.

##### TF2, TF12, TF88, TF682, TF763

Several OCS probe sets were synthesized for the purpose of enriching transcripts from different sets of human transcription factors (TFs). To make pools, ORF clones in the form of bacterial stocks were cherry-picked from the hORFeome collection using Genesis Automated Liquid Handler (Tecan). PCR success was checked by running a subset of the reactions on an E-Gel 96 Agarose Gel (Thermo Fisher Scientific). Pools corresponding to 2, 12, 88, and 682 ORF amplicons were used as template for Biotin-labeling PCR. For TF763, several pools were made, each containing ORFs of a similar length. Each pool underwent a separate Biotin-labeling PCR. TFs belonging to each probe set may be found in Supplementary Table 4.

### Sequence validation of OCS probes

Each OCS probe set was subjected to Illumina sequencing to verify the probe identities and abundances across the source templates. The Kapa DNA Hyper prep kit (Roche) was used, in which barcoded Illumina adapters were ligated directly to the probes. Samples were prepared on a Beckman Coulter Biomek FX. For each sample, approximately 50,000 paired-end reads of length 150 bp were generated on an Illumina MiSeq instrument.

To estimate the representation of each ORF within a given probe set, Kallisto^29^ (version 0.44.0, default parameters) was used to estimate gene-level abundances. Paired end reads were analyzed. Alignment indices were prepared from a FASTA file containing all human ORFeome sequences. For analysis of probe sets involving ERCC or SIRV standards, the relevant sequences were included in the FASTA file.

To generate read coverage plots across the ORF, reads were first aligned to the human ORFeome using Bowtie2^47^ (version 2.2.3) using “—local” option with default parameters. The alignment file, in SAM format, was parsed using SAMtools^48^ (version 1.2) and custom Python scripts were used to extract read coverage across the ORF on a per-nucleotide basis.

### Commercial probe synthesis

IDT Lockdown probes (Integrated DNA Technologies) were designed and synthesized for ARNTL, STAT1, and ZNF268 (Supplementary Table 1). The probes are high purity oligonucleotides (120-mers) with a biotin conjugated at the 5’ end. For each target ORF, a ∼1X tiling density was maintained by designing probes that randomly tile the forward and reverse complement sequences. Following reconstitution to 0.75 pmol/µl with TE buffer, probes from the targets were combined in equimolar ratios.

### Quantitative PCR (qPCR) method development

#### Preparation of standards

Standard solutions of ARNTL, STAT1, ZNF268, MBP, and HSP90AB1 were prepared for use in absolute quantification and qPCR method validation. ORF inserts were amplified from GATEWAY clones using M13 primers and Platinum Taq, as described in <u>Generating ORF amplicons</u>. Products were purified with 1.8X volume of Ampure XP beads and amplicons were run on an agarose gel to confirm presence of a single band of expected size. Final concentrations were measured by the Qubit dsDNA HS assay (Invitrogen) and Nanodrop spectrophotometry (Thermo Fisher Scientific). Molarity of each amplicon was calculated based on their sequence and concentration (e.g., ng/µl), accounting for vector backbone.

#### Assay development

TaqMan PCR assays (Integrated DNA Technologies) were designed against 450-500 bp regions within each of the five genes. The long target region length was necessary to specifically quantify full-length target cDNAs without background interference from the OCS or IDT oligonucleotide probes (data not shown). The PrimeTime Gene Expression Master Mix and accompanying protocol was used following manufacturer’s protocols, except for the extension time, which was increased to 120 seconds. Because of the unconventionally long qPCR target length, we performed full validation of the qPCR method and established excellent linearity (r^2^=1.00), precision (0.73-1.13 % CV), and limit of detection (3.2e-16 to 3.2e-10 M) for each of the five genes.

### Preparation of cDNA

cDNA was prepared using the SMARTer cDNA synthesis kit (Clontech). Approximately 1 µg of total RNA was input per reaction. All manufacturer’s protocol was followed, except for the use of a custom oligo(dT) containing a 16-mer barcode at the 5’ end, thereby uniquely labeling each cDNA preparation (Supplementary Table 12). After cDNA synthesis, whole cDNA amplification was performed so as to generate enough cDNA for capture reactions. The number of PCR cycles was optimized so as to avoid overamplification; this was done by monitoring product formation in small-scale PCR reactions and checking the product on agarose gels.

### Preparation of spike-in mixtures

#### ERCC spike-ins

Spike-in mixtures were prepared in which 1 µl of a 1:10, 1:80, or 1:5120 dilution of ERCC RNA Spike-In Mix (mix 1, Thermo Fisher) were each combined with 1 µg of UHRR. Each mixture was converted to cDNA as in the section **Preparation of cDNA**.

#### SIRV spike-ins

Spike-in mixtures were prepared in which 2.5 µl of a 1:10 dilution of SIRV RNA Spike-In Mix (mix E0, Lexogen) were each combined with 1 µg of UHRR. Each mixture was converted to cDNA as in the section **Preparation of cDNA**.

### Full-length cDNA enrichment

This protocol was adopted from the following two protocols: 1) “Hybridization capture of DNA libraries using xGen® Lockdown® Probes and Reagents” from IDT (version 2) and 2) “cDNA Capture Using IDT xGen® Lockdown® Probes” from PacBio (Part Number 101-604-300 Version 01).

#### Preparation of cDNA

Approximately 1 µg of purified cDNA was combined with 1 nmol of Clontech primer and 1 nmol of oligo(dT)_18_ containing a three-carbon spacer at the 3’ end (Eurofins Scientific), oligonucleotides that serve as blockers. The solution was dried down using vacuum centrifugation and subsequently resuspended in 8.5 µl 2X hybridization buffer, 2.7 µl enhancer buffer, and 1.8 µl of water, reagents supplied from the IDT Lockdown xGen kit.

#### Hybridization experiment

The cDNA was heated to 95C for 10 minutes, followed by a ramp down to 65C. Either 4 ng of OCS probes or 3 pmol of IDT probes were added and the mixture was incubated at 65C for 4 hours. 50uL of M-270 streptavidin beads (Invitrogen) were added and a series of washes were performed according to the IDT xGen Lockdown protocol version 2, except that initial washes used wash buffer pre-heated to 72C instead of 65C to reduce non-specific binding.

#### On-bead PCR

After the washes, the final bead solution was resuspended in 50 µl of TE buffer. To amplify the full-length cDNAs that were captured on the beads, on-bead PCR was performed with 5 µl of resuspended beads in a 30 µl reaction using KAPA HiFi HotStart 2X mix (KAPA) and the universal Clontech primer. The program was run for 30 cycles, with each cycle consisting of denaturation at 98°C for 20 seconds, annealing at 65°C for 15 seconds, and extension at 72°C for 5 minutes. The final extension was for 10 minutes.

#### Heat elution

Heat elution was performed for the purpose of quantifying abundances of bound cDNA via qPCR. An aliquot of beads was diluted 10-fold with buffer EB (10 mM Tris-HCl, pH 8.0) (Qiagen) and heated at 95C for 5 minutes. Beads were placed on a magnet and supernatant recovered for subsequent qPCR analysis.

### Enrichment of TFs from 7-tissue cDNA

A capture experiment was performed using TF763 against the pool of all seven cDNA tissues. The PacBio platforms have a slight length bias against longer cDNAs during loading. Therefore, to increase the recovery of transcripts across longer lengths, a second capture was performed to increase recovery of longer transcripts. The seven tissue cDNA was size selected using SPRISelect (Beckman Coulter) so that only transcripts above ∼2kb was recovered. A second capture was performed using TF763 against the 2kb+ size-selected cDNA. The capturant involving the full-size cDNA and the capturant involving the 2kb-size-selected cDNA were each sequenced on a 1M SMRTcell on the PacBio Sequel system. Therefore, a total of two 1M SMRTcells were run for the capturant. The original, unenriched input cDNA was also sequenced on an independent 1M SMRTcell.

### Illumina library preparation and analysis

#### Quantification of transcript abundances in brain

cDNA was synthesized from total RNA from the cortex region of human brain (Biochain) using the protocol described in **Preparation of cDNA**. cDNA was converted into an Illumina library using the NEBNext protocol (New England Biosciences) and ∼20 million PE75 reads were sequenced on an Illumina NextSeq, in duplicate. Sequencing data was collected at the Center for Cancer Computational Biology (CCCB) at the Dana-Farber Cancer Institute.

To estimate expression values for each gene, RSEM was used with the STAR aligner. The STAR genome index was built based on hg38 and using annotation obtained from GENCODE (version 27). Transcripts per million (TPM) values were calculated using RSEM (version 1.2.29)^49^.

#### Sequencing and quantification of enriched cDNA from capture experiments

The following sequencing data was collected for the enrichment of three TFs (Fig. 1d), ERCC ORFs (Fig. 2b), and the other TFs studied (Fig. 4a). Illumina sequencing data was collected following the procedure described in **Sequence validation of OCS probes**. To quantify isoform- and gene-level expression, Kallisto (version 0.44.0) was used using default parameters. To estimate values for all human genes (as in Fig. 1d, 4a), Kallisto indices based on GENCODE (version 27) transcript sequences were used. To estimate expression values for each ERCC ORF, Kallisto indices based on ERCC and GENCODE (version 27) sequences were used.

For the TF multiplexing experiment described in Figure 3, Illumina sequencing data was collected on cDNA subjected to a workflow similar to Nextera sequencing (Plexwell sequencing, SeqWell). For gene quantification, Kallisto (version 0.44.0) was used with default parameters.

#### Orthogonal validation of TF isoforms - RNA-seq

Human tissue total RNA samples were converted to Illumina libraries using the KAPA mRNA Hyper Prep kit, following manufacturer’s protocol (KAPA). Libraries were barcoding using TruSeq Illumina Adapters Sets A and B (Illumina).

### PacBio library preparation

For each reaction, ∼1 µg of either input cDNA or captured cDNA was converted into a SMRTbell library using the SMRTbell Template Prep Kit 1.0 (Pacific Biosciences) and sequenced on either a PacBio RSII or Sequel system (Pacific Biosciences).

### PacBio data analysis with Iso-Seq 3

Bioinformatics analysis was done by running the Iso-Seq 3 application in the PacBio SMRT Analysis v6.0 to obtain high-quality, full-length transcript sequences, followed by downstream analysis, as described below.

#### Identification of full-length reads

Full-length reads were determined as circular consensus sequence (CCS) reads that contained both the 5’ and 3’ primer and the polyA tail preceding the 3’ primer. The 5’ primer consists of the Clontech SMARTer cDNA primer with an ATGGG overhang. The 3’ primer consists of a 16-bp PacBio barcode that is sample-specific followed by the Clontech SMARTer cDNA primer.

#### Isoform-level clustering analysis to obtain high-quality transcript sequences

To increase detection of rare isoforms, the de-multiplexed full-length reads were pooled to perform isoform-level clustering analysis^35^. After clustering, consensus sequences were called using the Arrow algorithm and only polished sequences with predicted consensus accuracy ≥ 99% were considered high-quality and retained for the next step.

#### Mapping to hg38 and filtering for on-target isoforms

The high quality transcript sequences were mapped to hg38 using minimap2^50^ (version 2.11-r797) using parameters “-ax splice -t 30 -uf --secondary=no -C5”. We then filtered transcripts mapped to targeted probe region with ≥ 99% coverage and ≥ 95% identity.

#### De-multiplex final isoforms by sample

We recovered the relative abundance of each of the final isoforms in each sample by extracting the fraction of full-length reads supporting each isoform from each sample.

#### Isoform annotation and quality assessment with SQANTI2

SQANTI is a computational tool for annotation and quality assessment of full-length isoforms sequenced on long-read platforms^36^. We adapted SQANTI so that it includes additional functional and quality features relevant to isoform quality, a version called SQANTI2 (unpublished, github: https://github.com/Magdoll/SQANTI2). All de-multiplexed isoforms from the Iso-Seq 3 pipeline was processed with SQANTI2 using default parameters. Isoform and junction annotation and feature files, including match information to GENCODE version 29, were output.

### Isoform overlap estimation

GENCODE v29 GTF-based annotations were parsed to determine for each protein-coding gene the Appris-annotated “reference” isoform. The exon-intron structures for each Appris isoform was compared with every isoform (with a “basic” tag) of that gene using bedtools intersect function (v2.27.1). SIRV isoform overlap was calculated using a similar routine.

### GENCODE recoveries

For each GENCODE isoform (i.e., ENST) in version 29, we determined if there was an exact match in the PacBio transcripts. An exact match is defined as cases in which the detected full-length isoform contains an exact sequence of junctions (i.e., introns) as found in the GENCODE transcript. This was accomplished using a modified version of the SQANTI program, SQANTI2.

### Functional features of isoforms annotated within SQANTI2

#### CAGE peak overlap

CAGE peak annotations^51^ were downloaded from:

> http://fantom.gsc.riken.jp/5/datafiles/latest/extra/CAGE_peaks/hg19.cage_peak_phase1and2combined_coord.bed.gz

Genomic coordinates were converted from hg19 to hg38 using the liftOver program from the UCSC Genome Browser^52^. The genomic position of the 5’ end of the isoforms was compared to all CAGE peaks and the following criteria was determined: 1) the distance between the 5’ end and the center position of the closest CAGE peaks, and 2) whether the 5’ resided within a range of a CAGE peak.

#### Junction conservation

The conservation at each junction was obtained through downloading phyloP scores for each nucleotide in the human genome (hg38)^53^. PhyloP conservation scores for each donor and acceptor were obtained. The dinucleotides at the splice donor (e.g., GT) as well as the adjacent nucleotide residing on the exon was analyzed for the 5’ splice site. The dinucleotides at the splice acceptor (e.g., AG) as well as the adjacent nucleotide residing on the exon was analyzed for the 3’ splice site. Therefore, a trinucleotide was analyzed for each splice site.

#### polyA motif

A polyA motif is commonly found upstream of the site of cleavage and polyadenylation. The highest frequency polyA motifs in human are AAUAAA and AUUAAA, considered canonical motifs due to their high frequency^54^. The genomic position of the 3’ end of the isoforms was located, and it was determined whether there was presence of a canonical motif 5-25 nucleotides upstream of the 3’ site.

## Acknowledgements

We thank members of the Center for Cancer Systems Biology (CCSB) at the Dana-Farber Cancer Institute (DFCI) for helpful comments and discussion, Zach Herbert from the DFCI Molecular Biology Core Facilities, Shana McDevitt from the Vincent J. Coates Genomics Sequencing Laboratory at the California Institute for Quantitative Biosciences (QB3) at the University of California, Berkeley, Stuart Levine at the MIT BioMicro Center for help with sequencing, and Chris Burkhart for technical assistance. SIRV amplicons were a gift from Lexogen GmbH, Vienna, Austria. This work was supported by an NHGRI Center of Excellence in Genomic Science grant P50HG004233 and an NCI Cancer Systems Biology Consortium grant U01CA232161 awarded to M.V. G.M.S. was supported by NIH training grant T32CA009361, a Charles A. King Trust Postdoctoral Research Fellowship, and a Melanoma Research Foundation Career Development Award.

## Author contributions

G.M.S. conceived of the project through ongoing discussions with and feedback from J.G.U., M.L.S., R.S., D.E.H., M.A.C. and M.V. G.M.S. designed experiments, under the guidance of D.E.H. and M.V. G.M.S. and K.T. performed experiments. G.M.S., E.T., and T.H. analyzed the data. L.Y. and D.D. analyzed Illumina data of probe sequences. G.M.S. wrote the manuscript, with contributions from all other authors.

## Additional information

**Correspondence and requests for materials** should be addressed to G.M.S. and D.E.H.

## Supplementary information

**Supplementary Fig. 1.**
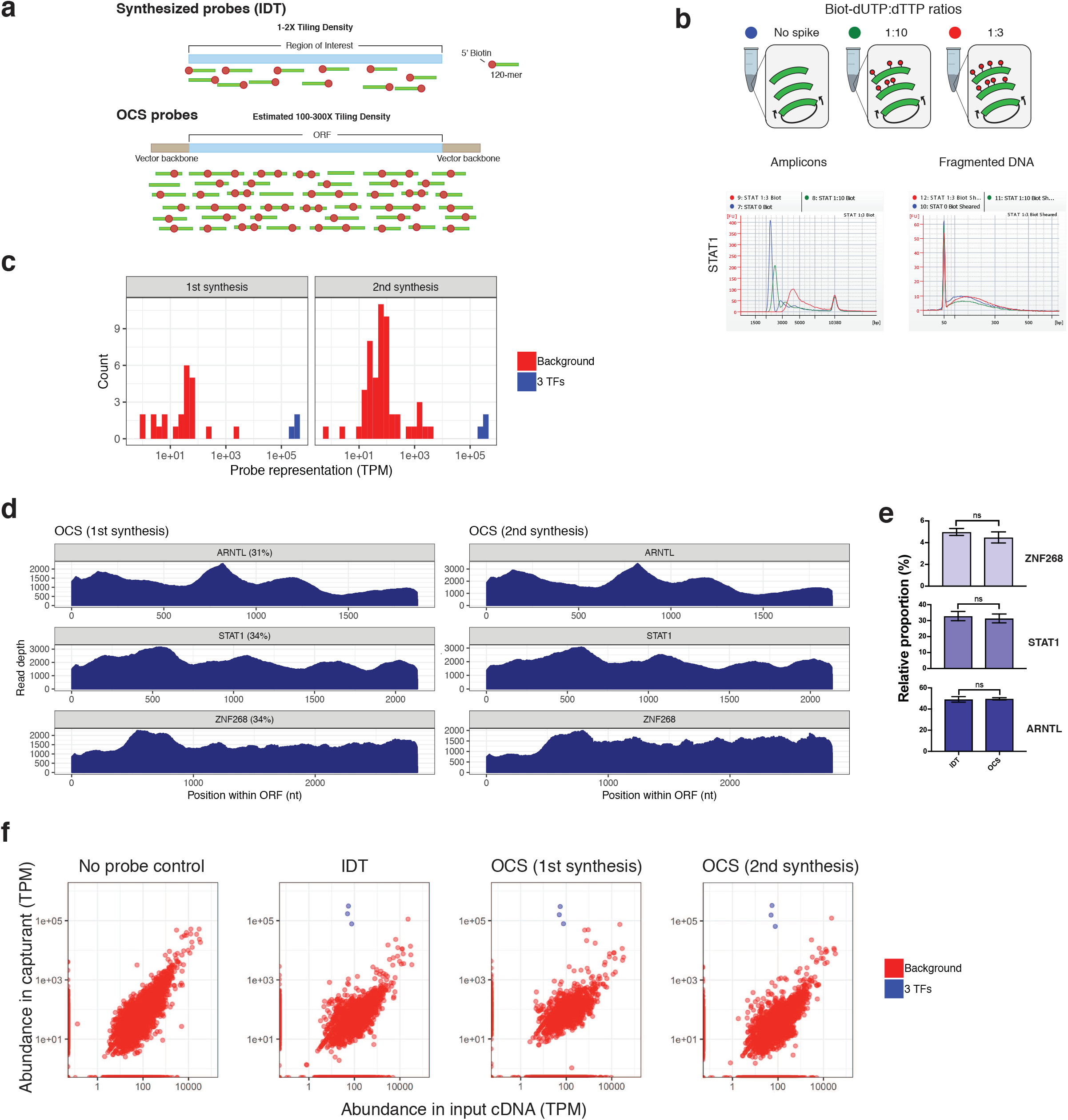
OCS and IDT probe sets perform comparably. **a** Schematic of commercially synthesized (IDT) probes and OCS probes. **b** QC of Biotin-dUTP-labeling PCR and fragmentation. Biotin-dUTP replaced dTTP in the PCR mix at ratios of 1:10 and 1:3. The lower panel shows Bioanalyzer traces of the amplicons before and after sonication (Methods). Color of legend dots and plotted lines (blue, green, red) correspond to different levels of Biotin-dUTP spikes. **c** Purity of probe sets. Abundance of probes from on-target (dark blue) versus off-target (red) ORFs. **d** Probe coverage across the source ORF templates. Read depth is the number of aligned reads at each nucleotide position. Reads were sequenced on a MiSeq (Methods). **e** Comparison of on-target proportions between IDT and OCS probes. P-values were calculated using the Mann-Whitney two-tailed test. Each replicate represents an independently executed capture reaction. Error bars, s.e.m. (IDT, n=6; OCS, n=9); n.s., not significant (P-value is above 0.05). **f** Background binding profiles. Comparison of gene abundance of input and capturant for each experiment. TPM, transcripts per million.

**Supplementary Fig. 2.**
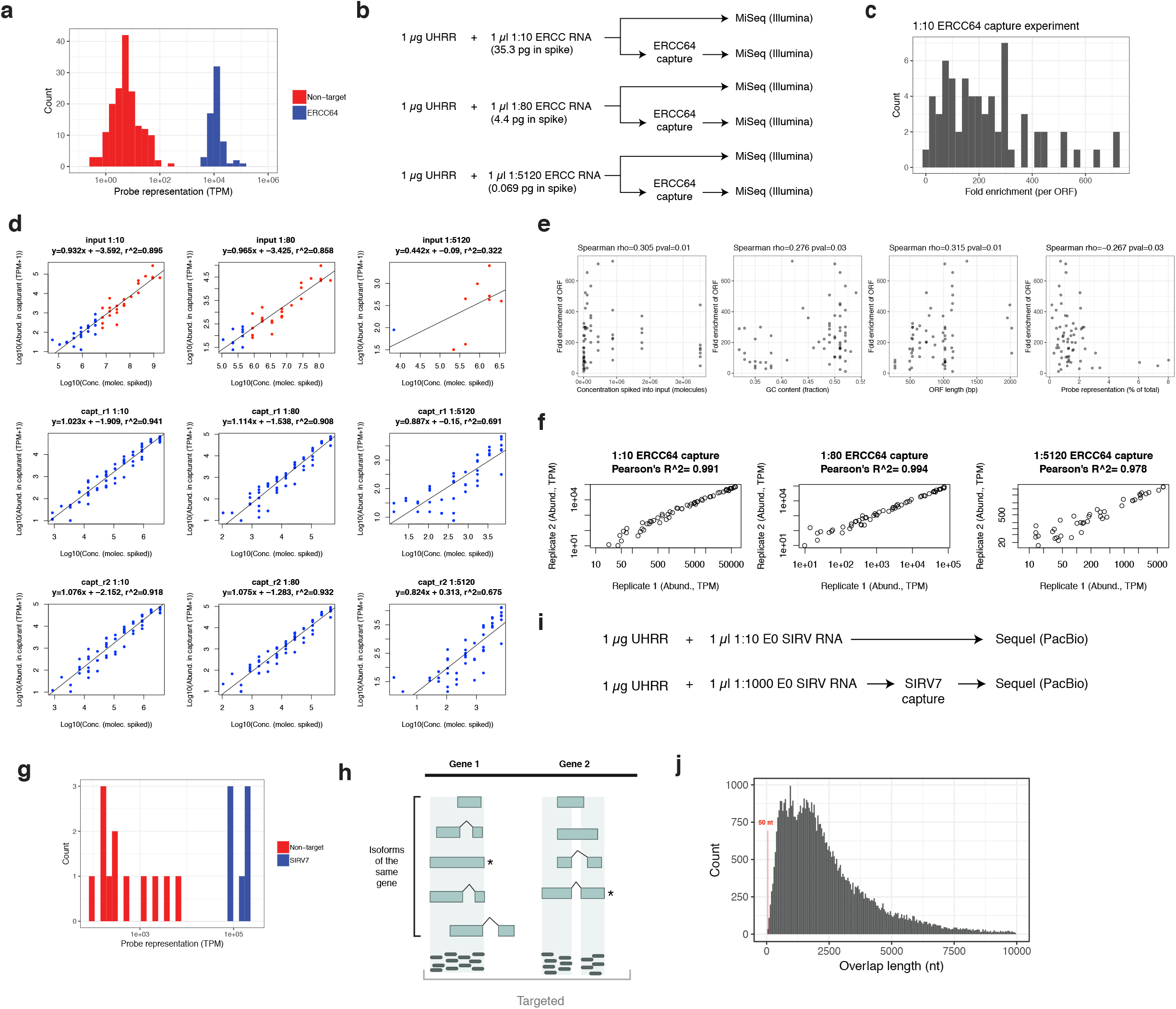
Benchmarking OCS analytical performance with spike-in standards. **a** Purity of ERCC64 probe set. Plots show abundance of probes derived from targeted ERCC templates (blue) versus non-targeted genes (red). **b** Schematic of ERCC spike-in capture experiment. 1:10, 1:80, and 1:5120 are 10-, 80-, and 5120-fold dilutions of the ERCC spike-in mix 1. ERCC, External RNA Controls Consortium; UHRR, universal human RNA reference. **c** Distribution of ERCC ORF-specific fold enrichments observed for the experiment involving the 1:10 spike-in. **d** Linearity of ERCC standards, before and after capture. Linear regression performed on all 92 ERCC ORFs for input (top panel), and 64 targeted ERCC ORFs in the capturants (middle and bottom panels). Data for two independent captures (r1, r2) are shown. Equation of best fit line and R^2^ is shown for each plot. **e** Plots showing relationship between ERCC ORF properties and enrichment efficiency. Spearman’s rho and associated p-value shown. **f** Reproducibility of technical replicates of ERCC64 captures. Pearson’s correlation calculated for the 64 ERCC ORFs. **g** Purity of SIRV7 probe set. Plots show abundance of probes derived from targeted SIRV templates (blue) versus non-targeted genes (red). Note that one SIRV per locus was selected to be included in the probe set. **h** Schematic of the SIRV experiment. Isoforms with an asterisk mark the representative SIRV selected for each locus. **i** Schematic of the SIRV capture experiment. SIRV, Spike-in RNA Variant Control; UHRR, universal human RNA reference. **j** Distribution of overlap lengths between the GENCODE principal isoform and all isoforms of that gene. GENCODE version 29 was used in this analysis. The position of 50 nt is marked by the red line, denoting the threshold under which isoform enrichment efficiency is expected to marked decrease.

**Supplementary Fig. 3.**
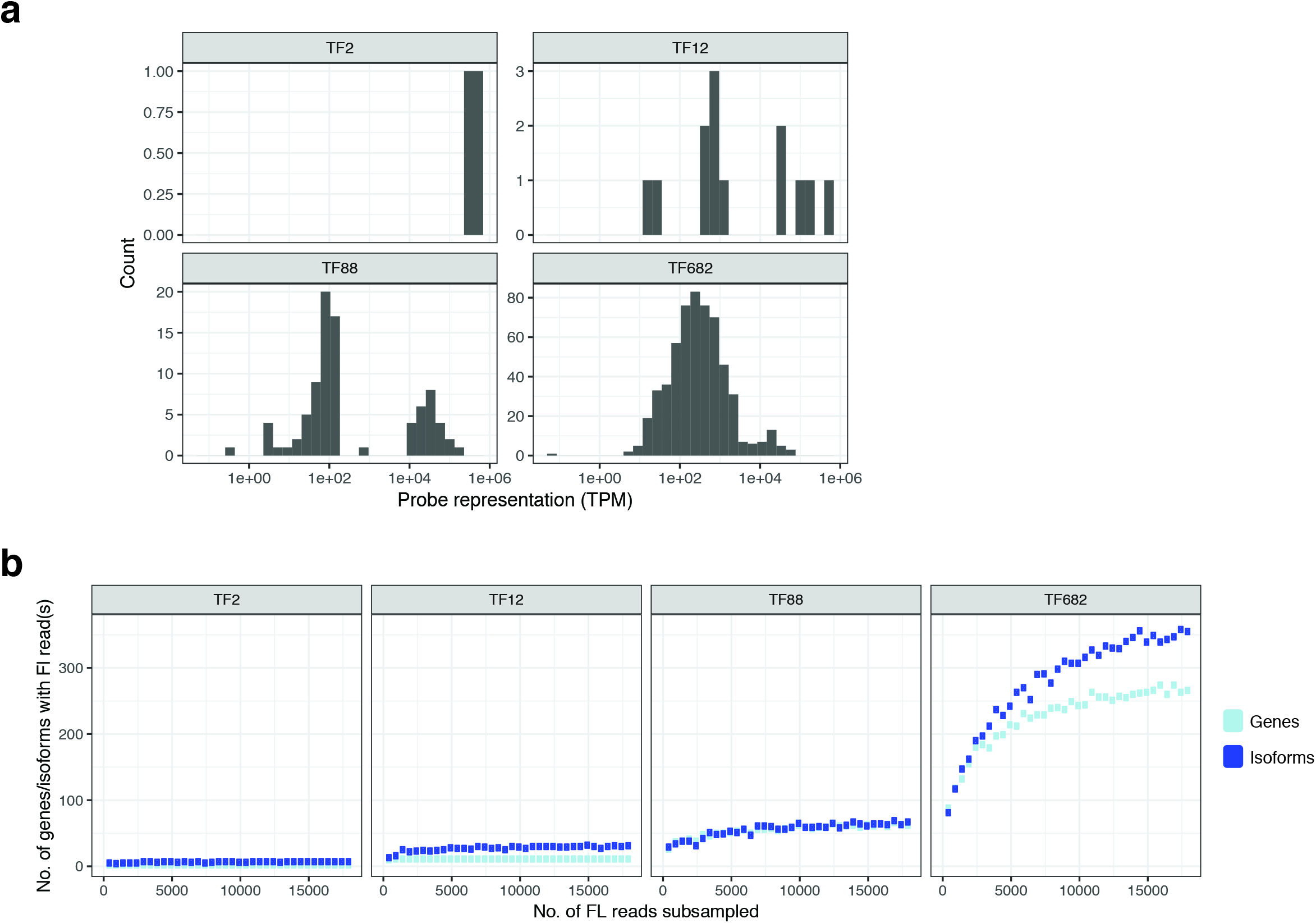
Multiplexing OCS captures. **a** Distribution of the abundances of probes, on a per-ORF-basis. Results for TF2, TF12, TF88, and TF682 are shown. X-axis shows transcripts per million, which was calculated per ORF. **b** Saturation-discovery curves for number of detected genes (dark blue) and isoforms (light blue). Full-length reads were subsampled without replacement from the original data and number of genes or known isoforms recovered recalculated.

**Supplementary Fig. 4.**
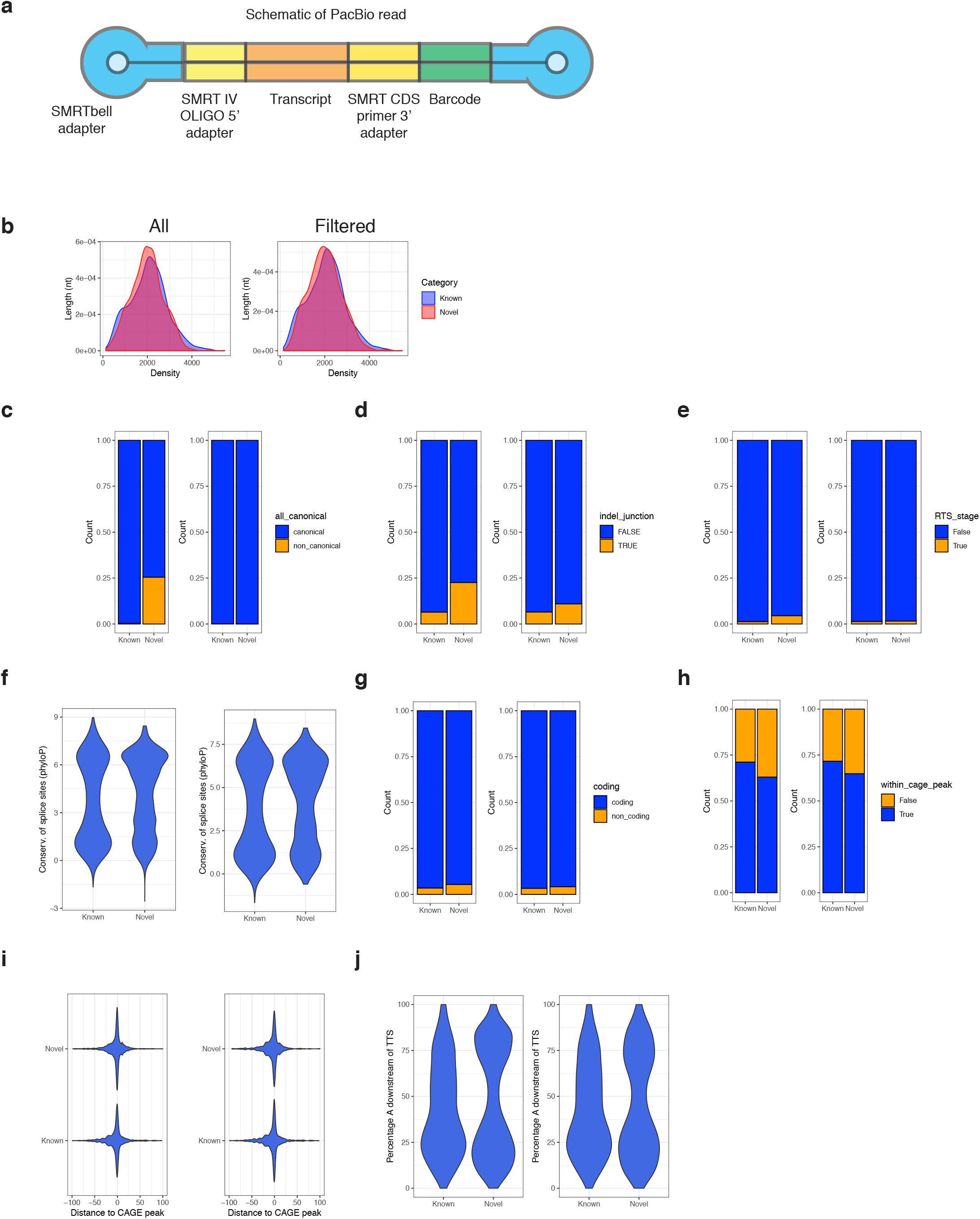
Enrichment and characterization of novel TF isoforms. **a** Depiction of structure of final PacBio reads after library preparation. **b-j** Comparison of sequence and functional features between known and novel isoforms, before and after filtering based on Illumina data. Comparisons include **b** distribution of the transcript length; proportion of isoforms containing **c** non-canonical junctions, **d** indel sequencing errors adjacent to the splice junction, and **e** a predicted reverse transcription template switching artifact; **f** distribution of phyloP-based conservation at nucleotides residing adjacent to the junction (Methods); proportion of isoforms which **g** are predicted as containing a coding ORF (SQANTI, GMST, see Methods), and **h** contain a 5’ end residing within a CAGE peak (Methods); **i** distribution of distances between 5’ end of isoform and an annotated CAGE peak, and **j** distribution of the percentage of A/T content, on the genome, which is immediately downstream of the 3’ site as detected by the sequenced isoform.

## Tables

Supplementary Table 1 – Sequences of biotinylated oligos synthesized for capture of TFs in Figure 1.

Supplementary Table 2 – List of ERCC ORFs belonging to probe set “ERCC64”.

Supplementary Table 3 – Primers used to amplify SIRVs for probe synthesis.

Supplementary Table 4 – List of genes targeted in the TF multiplexing experiment. Includes the ORF sequences used as template for probe synthesis.

Supplementary Table 5 – Abundance of probes within each probe set in the TF multiplexing experiment. Units are in transcripts per million (TPM).

Supplementary Table 6 – Variability in probe abundance within each probe set in the TF multiplexing experiment. Summary statistics are based on distribution ORF-specific TPMs.

Supplementary Table 7 – List of genes targeted in the TF enrichment experiment.

Supplementary Table 8 – Abundance of probes within each probe set in the TF enrichment experiment. Units are in transcripts per million (TPM).

Supplementary Table 9 – Proportion of GENCODE genes and isoforms detected in the TF enrichment experiment.

Supplementary Table 10 – GENCODE isoforms, categorized by their transcript support levels (TSLs), detected in the TF enrichment experiment.

Supplementary Table 11 – High-quality isoform sequences detected in the TF enrichment experiment. Match category, as defined within the SQANTI program, is listed for each isoform.

Supplementary Table 12 – Oligo(dT) barcode sequences used in first strand synthesis during cDNA preparation.

